# Piscichuviral encephalitis in marine and freshwater chelonians: first evidence of jingchuviral disease

**DOI:** 10.1101/2023.02.24.528524

**Authors:** Weerapong Laovechprasit, Kelsey T. Young, Brian A. Stacy, Steven B. Tillis, Robert J. Ossiboff, Jordan A. Vann, Kuttichantran Subramaniam, Dalen Agnew, Jian Zhang, Shayna Whitaker, Alicia Walker, Andrew M. Orgill, Lyndsey N. Howell, Donna J. Shaver, James B. Stanton

**Affiliations:** Department of Pathology, College of Veterinary Medicine, University of Georgia, Athens, GA; Office of Protected Resources, National Marine Fisheries Service, National Oceanic and Atmospheric Administration, University of Florida, Gainesville, FL and Pascagoula, MS; Department of Comparative, Diagnostic, and Population Medicine, College of Veterinary Medicine, University of Florida, Gainesville, FL; Department of Infectious Diseases and Immunology, College of Veterinary Medicine, University of Florida, Gainesville, FL; Emerging Pathogens Institute, University of Florida, Gainesville, FL; Department of Pathobiology and Diagnostic Investigation, College of Veterinary Medicine, Michigan State University, Lansing, MI; Amos Rehabilitation Keep at University of Texas Marine Science Institute, Port Aransas, TX; National Park Service, Padre Island National Seashore, Corpus Christi, TX

**Keywords:** chuvirus, alligator snapping turtle, kemp’s ridley turtle, chuviridae, *Lepidochelys*, *Macrochelys*, chelonian, MinION, deep sequencing

## Abstract

Chuviruses (family *Chuviridae*), which are in the recently discovered order *Jingchuvirales*, were first identified in arthropods in 2015 and have been detected through metagenomics in numerous invertebrates, but only a few vertebrates. With only few metagenomically based detections in vertebrates, their replication competency in vertebrates remained questioned, let alone their pathological significance. This study identified three novel chuviruses as the etiology of lymphocytic meningoencephalomyelitis in three wild aquatic turtles: an alligator snapping turtle (*Macrochelys* sp.), a Kemp’s ridley turtle (*Lepidochelys kempii*), and a loggerhead turtle (*Caretta caretta*). The application of random, deep sequencing successfully assembled the complete snapping turtle chuvirus-1 [STCV-1], Kemp’s ridley turtle chuvirus-1 [KTCV-1] genome, and loggerhead turtle chuvirus-1 [LTCV-1]) genome. The STCV-1 and KTCV-1 sequences were used to create custom RNAscope^™^ probes for *in situ* hybridization, which confirmed STCV-1, KTCV-1, and LTCV-1 (cross reactivity to the KTCV-1 probe) RNA within the inflamed region of the brain and spinal cord. STCV-1 and KTCV-1 were isolated on several turtle-origin cell lines. Phylogenetic analysis illustrated that all three viruses clustered with other vertebrate chuviruses, all within the genus *Piscichuvirus*. With more than 91% pairwise amino acid identity of the polymerase proteins, STCV-1, KTCV-1, and LTCV-1 belong to the same novel species, putatively named Piscichuvirus testudinae. This study demonstrates the first *in situ* evidence of chuviral pathogenicity in animals and only the second instance of jingchuviral isolation. The association of these chuviruses in three different chelonians with neurologic disease suggests the possibility that chuviruses are a significant, previously unrecognized cause of lymphocytic meningoencephalomyelitis in freshwater and marine turtles. Additional studies of these viruses are needed to elucidate their role in chelonians and the possibility of related viruses in other related hosts.

**Importance:** Chuviruses have been identified in multiple animal species, including humans. However, most were identified metagenomically, and detection was not strongly associated with disease. This study provides the first evidence of chuviral disease in animals in diseased tissue: mononuclear meningoencephalomyelitis in three chelonians from three different genera, two distinct families. These pathogenic turtle chuviruses belong to the genus *Piscichuvirus* containing other non-mammalian vertebrate chuviruses and were classified together within a novel chuviral species. This study supports the need for further investigations into chuviruses to understand their biology, pathogenic potential, and their association with central nervous system inflammation in chelonians, other reptiles, and other vertebrates.

## Introduction

Mononuclear meningoencephalomyelitis (inflammation of the meninges, brain, and spinal cord) characterized by perivascular cuffing often warrants a differential diagnosis of viral infection. Determining the etiology of such diseases can be challenging in any species, but is particularly so in non-domesticated animals due to limited diagnostic assays and knowledge gaps regarding infectious agents of understudied taxa (1–3). In mammals, a number of viruses are associated with this histopathologic presentation, including herpesviruses (4), alphavirus (5), flaviviruses (6), morbilliviruses (7), etc. In reptiles, infection with herpesviruses (8), adenoviruses (9, 10), picornaviruses (11), arenaviruses (12), and paramyxoviruses (13) are potential causes of mononuclear meningoencephalomyelitis. While spirorchiids (14) and mycobacteria (15) can infect the central nervous system of sea turtles, the inflammation is more lymphohistiocytic to granulomatous. However, a large proportion of suspected viral encephalitis cases remain idiopathic, which prevents understanding the true impact on animal populations and collections (4, 16). Random sequencing has provided an unbiased mechanism to detect viruses, resulting in the identification of arenavirus encephalitis in human (17), astrovirus encephalitis in cattle (18), and the recently discovered turtle fraservirus-1 encephalitis in freshwater turtles (19).

Metagenomic surveillance of RNA viruses in various invertebrate species recently led to the discovery of chuviruses (20), which are negative-sense, single-stranded RNA viruses in the family *Chuviridae*, order *Jingchuvirales* (21). To date, 48 chuviral species have been characterized and taxonomically reclassified by the International Committee on Taxonomy of Viruses (ICTV) (21). Chuviruses have since been detected in other phyla, e.g., platyhelminths, cnidaria, nematodes, mollusks, arthropods, echinoderms, and a few vertebrates (20, 22–28). Moreover, chuviral coding sequences have also been observed within BEL-Pao retrotransposons of several host species, such as insects and vertebrates (20). Therefore, chuviruses are also recognized as endogenous viral elements (EVEs) (29). This suggests that even though chuviruses are recently identified, they have circulated and co-evolved with animals along with other commonly recognized viruses. In fact, the varying genome organization of chuviruses (circular vs. linear and segmented vs. unsegmented) and the phylogenetic analysis of the RNA-dependent RNA polymerase suggest that these viruses have a unique history among the viruses in the class *Monjiviricetes*(20, 21).

In vertebrates, only five chuviruses have been identified: 1) hardyhead chuvirus (host: unspecked hardyhead *[Craterocephalus fulvus]*) (28), 2) Wēnlǐng fish chu-like virus (host: long-spine snipefish *[Macroramphosus scolopax]*) (25), 3) Guangdong red-banded snake chuvirus-like virus (host: red-banded snake *[Lycodon rufozonatus]*) (25), 4) Herr Frank virus 1 (host: boa constrictor *[Boa constrictor]*) (22), and 5) Nuomin virus (host: human) (30), all of which were detected through metagenomic sequencing. However, only two of these viruses have been identified in clinically ill patients. Herr Frank virus-1 was identified from three different snakes that were individually diagnosed with varying combinations of ulcerative stomatitis, osteomyelitis, and fibrino-necrotizing cloacitis, but these animals also had coinfections with other potentially pathogenic viruses (reptarenavirus and hartmanivirus) and no *in situ* localization of chuviruses was attempted (22), preventing attribution of the clinical signs and lesions to the chuvirus. Nuomin virus was identified from the serum of febrile human patients in China; however, no tissue-based studies were performed to support it as the cause of the fever (30) or to determine the pathology associated with infection. While these studies demonstrate the potential of chuviruses to infect vertebrates, no studies have demonstrated chuviruses in lesions in clinically ill animals.

The goal of this study was to determine the cause of idiopathic mononuclear meningoencephalomyelitis in three free-ranging aquatic turtles (alligator snapping turtle *[Macrochelys* sp.], Kemp’s ridley turtle *[Lepidochelys kempii]*, and loggerhead turtle *[Caretta caretta*]). Based on histological findings from all animals, it was hypothesized that unknown viruses were causing the mononuclear inflammation in the central nervous system, these viruses could be identified by random sequencing, and that viral RNA could be localized to diseased tissue supporting the causality of the viruses.

## Results

### Animal history and pathology

An adult male alligator snapping turtle (56.5 cm straight carapace length [SCL] from nuchal notch to caudal tip) was found minimally responsive on the shore of Lake Wauburg (29.524879°N; −82.300594°W) in Alachua County, Florida, USA and was brought to the University of Florida, College of Veterinary Medicine (UF-CVM) for care on June 10, 2009. As the location of discovery is outside of the species range, it was strongly suspected that this turtle was captured elsewhere and released. The animal was weak and lethargic upon evaluation and 5 days later was euthanized due to declining condition. The turtle was in good nutritional condition as evidenced by abundant body fat. Necropsy findings included a locally extensive ulcerated wound on the ventral tail base, infestation by small numbers of leeches, a single pentastome within the left lung, small numbers of unidentified nematodes within the large intestine, a chronic fracture of the left fibula with callus formation, and chronic arthritis of the left mandibular joint with erosion of the articular cartilages and remodeling of bone. The brain was grossly normal. Histopathological examination of the nervous system revealed mononuclear inflammation of the meninges, brain, and spinal cord (meningoencephalomyelitis). Moderate numbers of lymphocytes and plasma cells diffusely infiltrated the leptomeninx, extending into the adjacent brain and cranial nerves. All regions of the brain (including the olfactory bulbs, cerebrum, optic tectum, midbrain, cerebellum, and brainstem) (Fig. 1a) contained frequent lymphoplasmacytic cuffs. The associated neuroparenchyma was vacuolated and some neurons exhibited central chromatolysis (Fig. 1c). Similar inflammation disrupted the cervical spinal cord with lesions most concentrated within the gray matter. A viral etiology was considered most likely; brain tissue was negative for herpesvirus through conventional PCR (31).

**Figure 1.**
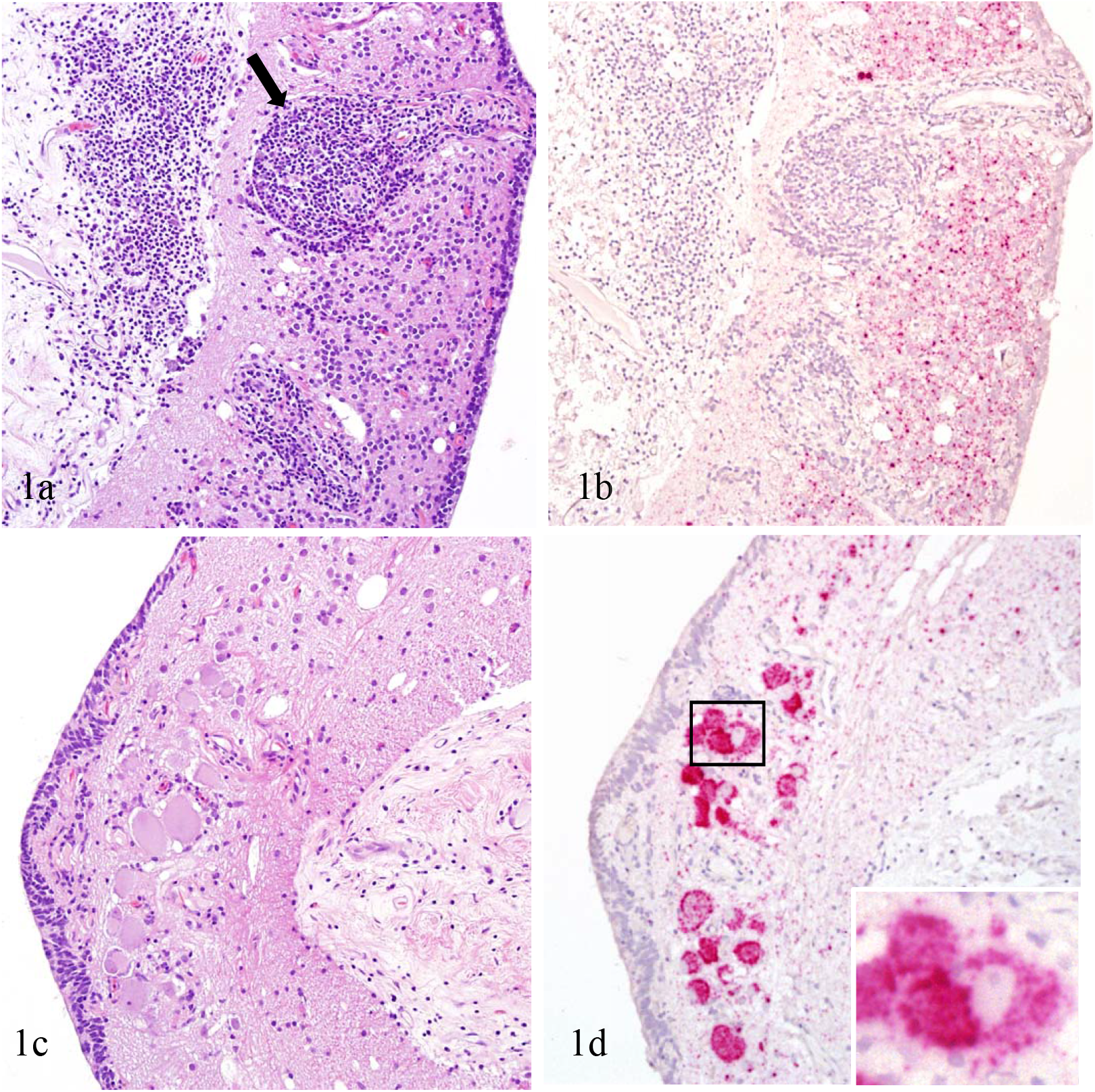

The second case was a subadult female Kemp’s ridley turtle (60.3 cm SCL) that was found stranded on Mustang Island Gulf Beach (27.67338°N; −97.16880°W) in Nueces County, Texas, USA and admitted to Amos Rehabilitation Keep (ARK) in Port Aransas, Texas. The animal was underweight, had accumulated epibiota (*Lepas* sp.) compatible with prolonged floating, and exhibited persistent neurological signs: circling to the left and right-sided asymmetric buoyancy. Euthanasia was elected due to quality of life concerns after 4.5 months of attempted treatment using antimicrobial, antiparasitic, and corticosteroid medications without clinical improvement. No gross abnormalities were observed in the nervous system or inner ears. Cytological evaluation of a postmortem cerebrospinal fluid (CSF) sample revealed marked histiocytic pleocytosis with a mild lymphocytic component. However, by histopathology the inflammation was predominantly lymphocytic and distributed as a severe, diffuse meningoencephalomyelitis with prominent perivascular cuffs and infiltration of cranial nerves (Fig. 2a). Neurons frequently contained eosinophilic, intranuclear inclusions and there was patchy vacuolation of the neuroparenchyma (Fig. 2a and 2c). There were no acid-fast organisms in sections of brain and a CSF slide stained using the Ziehl-Neelsen method. A viral etiology was suspected to be the cause of the lesion.

**Figure 2.**
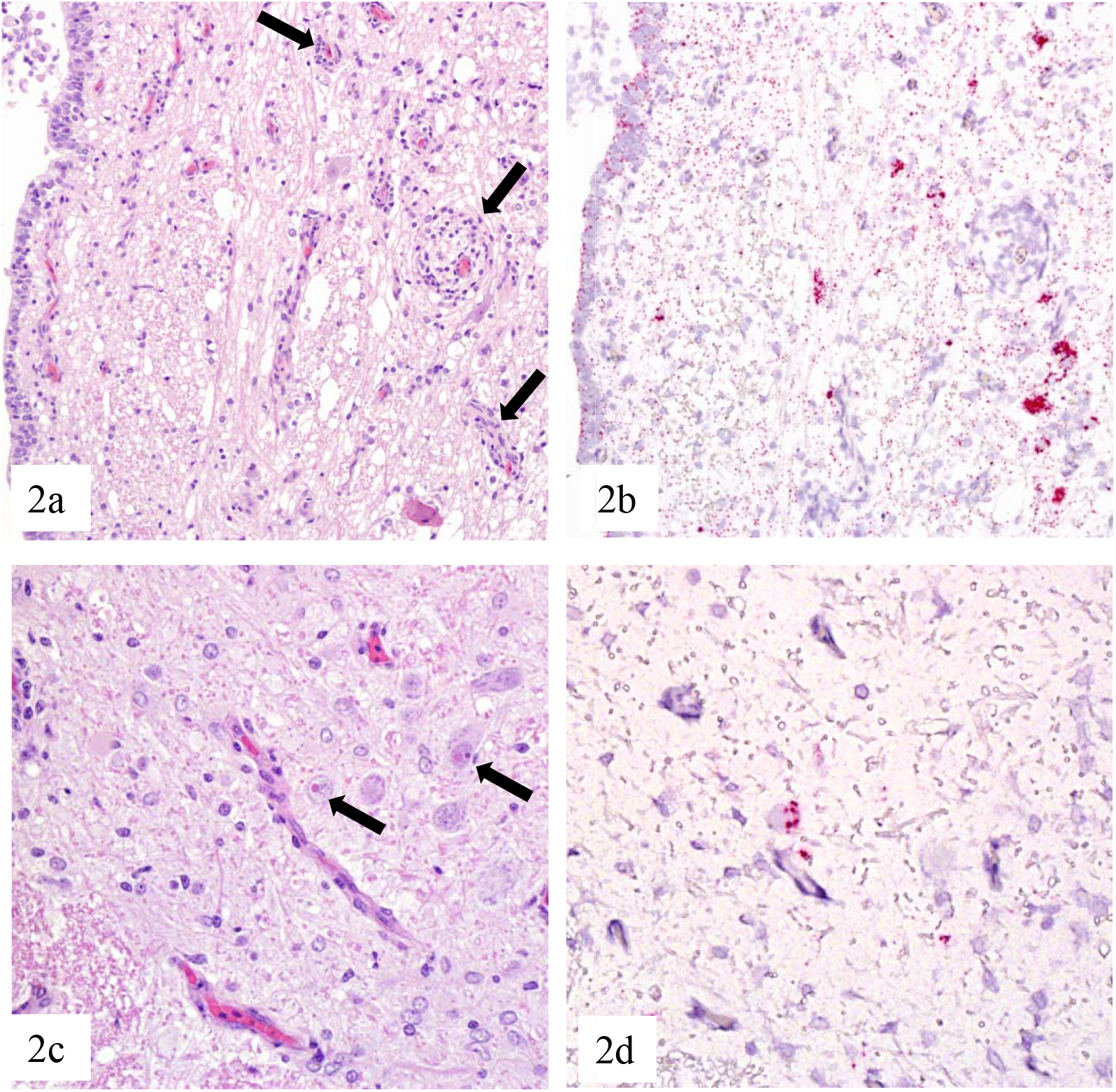

The third case was a stranded adult male loggerhead turtle (94.0 cm SCL) that was found unresponsive on a Gulf of Mexico beach (30.230591°N; −87.910237°W) in Baldwin County, Alabama, USA. The animal died soon after discovery. Nutritional condition was within normal limits based on the condition of skeletal muscle and abundance of body fat. Small (0.1–1.0 cm diameter) acorn barnacles suggestive of reduced activity were accumulated on the head, appendages, and shell. The central nervous system was grossly normal; however, significant inflammation was detected by histopathology. Moderate numbers of lymphocytes infiltrated the leptomeninx, neuroparenchyma, and cranial nerves, forming prominent perivascular cuffs (Fig. 3a and 3c). The inflammation was diffuse, but variable in intensity with relative severity in areas of the cerebrum, optic tectum, and cerebellum (Fig. 3a and 3c). There were no acid-fast organisms in sections of brain. A viral etiology was suspected to be the cause of the lesion.

**Figure 3.**
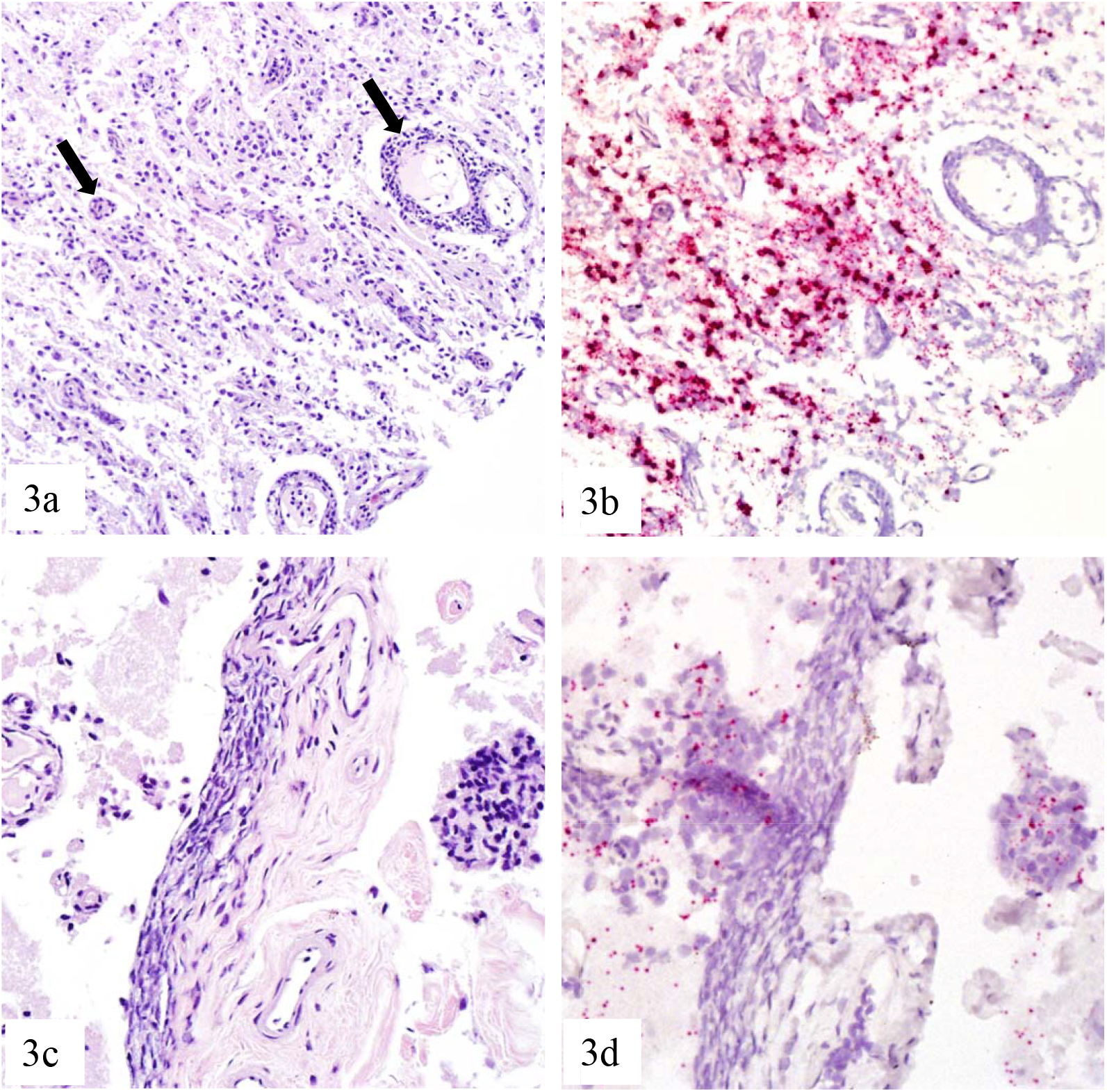

### Identification and characterization of viral genomes

#### Snapping turtle chuvirus (STCV-1)

Chuviral reads from MinION sequencing were initially detected through reference-based alignment using BLASTn. Out of 126,221 total reads, only 5 reads best aligned to a chuvirus (Wēnlǐng fish chu-like virus and Lampyris noctiluca chuvirus-like virus 1). Suspecting that the paucity of known chuviral sequences and the large sequence diversity of known chuviruses could lead to poor alignments, the reads were further interrogated through a custom index using a long-read aligner (i.e., Centrifuge, see Methods), resulting in thirty-one reads that best aligned to a chuvirus, including Wēnlǐng fish chu-like virus, Lampyris noctiluca chuvirus-like virus 1, and Herr Frank virus 1.

To assemble the genome, all reads from the sample that did not align via Centrifuge to the green turtle genome or to bacteria, and were larger than 50 nt run were mapped to Wēnlǐng fish chu-like virus in Geneious. This mapping resulted in the detection of 1,491 reads that built a complete genome for snapping turtle chuvirus-1 (STCV-1) with at least 10× coverage. The complete genome (10,781 nt) contains the complete CDS for the 5’-polymerase (*L*), glycoprotein (*G*), nucleoprotein (*N*), and viral protein-4 (*VP4)-3’* genes of the STCV-1. The *L, G, N*, and *VP4* genes were 6,438 nt, 2,052 nt, 1,446 nt, and 318 nt, respectively. The genomic structure of this novel chuvirus was linear and non-segmented, similar to other piscichuviruses. The 5’ untranslated region (UTR) was 91 nt and the 3’ UTR was 89 nt. In addition, 16 terminal nucleotides of each UTR were inverted repeats, with 3 nucleotide mismatches.

#### Kemp’s ridley turtle chuvirus (KTCV-1)

Raw reads were generated through random, deep Illumina sequencing. The BLASTX search of the assembly identified four contigs with significant identity to Herr Frank virus 1 (GenBank accession number MN567051) and Guangdong red-banded snake chuvirus-like virus (GenBank accession number MG600009), which were then used as a reference to create a draft genome of Kemp’s ridley turtle chuvirus-1 (KTCV-1). The complete genome contained 10,839 bases with four predicted coding regions: 5’-polymerase (*L*)-glycoprotein (*G*)-nucleoprotein (*N*)-viral protein-4 (*VP4*) −3’ of 6,438 nt, 2,052 nt, 1,500 nt, and 318 nt respectively. The genomic structure of this novel chuvirus was also linear and non-segmented. The 5’ UTR was 91 nt, and the 3’ UTR was 93 nt. In addition, 16 terminal nucleotides of each UTR were inverted repeats with 2 nucleotide mismatches.

#### Loggerhead turtle chuvirus (LTCV-1)

All 1,839,435 reads from the random and targeted MinION sequencing that did not align via Centrifuge to the green turtle genome or to bacteria, and were larger than 50 nt run were mapped to Kemp’s ridley turtle chuviral consensus sequence in Geneious. This resulted in the detection of 258 reads that built a complete genome for loggerhead turtle chuvirus-1 (LTCV-1) with at least 10× coverage except the first 9 bases of 5’terminus, which had 6–9× coverage. The complete genome contained 10,839 bases with four predicted coding regions: 5’-polymerase (*L*)-glycoprotein (*G*)-nucleoprotein (*N*)-viral protein-4 (*VP4*) −3’ of 6,438 nt, 2,052 nt, 1,500 nt, and 318 nt respectively. The genomic structure of this novel chuvirus was also linear and non-segmented. The 5’ UTR was 91 nt, and the 3’ UTR was 93 nt. In addition, 16 terminal nucleotides of each UTR were inverted repeats with 2 nucleotide mismatches. The termini were 100% identical to the termini of KTCV-1.

#### Genome comparison of piscichuviruses

According to the recent taxonomical framework for the order *Jingchuvirales* (21), L amino acid identity is required for the characterization of the jingchuviral taxonomy. The percent pairwise amino acid identities <90%, <31%, and <21% support differentiation of jingchuviruses as a novel species, genus, and family, respectively (21). Percent pairwise amino acid identity analysis of the STCV-1, KTCV-1, and LTCV-1 revealed that the pairwise identity of the L protein between two marine chelonian chuviruses (e.g., KTCV-1 and LTCV-1) and the freshwater chelonian chuvirus was approximately 91.6%; whereas the pairwise identity between KTCV-1 and LTCV-1 was 99.8%. This percent identity is markedly different from the percent amino acid identities for the L protein when comparing these chelonian chuviruses to non-chelonian chuviruses (46.9–47.3% to Herr Frank virus 1; Table 1 and supplemental data). Based on these pairwise amino acid identities, all three chelonian chuviruses belong to the genus *Piscichuvirus*, and are grouped together within the same novel species, putatively named Piscichuvirus testudinae.

**Table 1:** Percent pairwise identities of predicted L amino acid sequences of viruses in *Piscichuvirus* genus

|  | LTCV-1* | KTCV-1 | STCV-1 | MN567051 | MG600009 | MG600010 | MW645030 | KX884439 |
| --- | --- | --- | --- | --- | --- | --- | --- | --- |
| Loggerhead turtle chuvirus |  |  |  |  |  |  |  |  |
| Kemp's Ridley turtle chuvirus | 99.442 |  |  |  |  |  |  |  |
| Snapping turtle chuvirus | 91.566 | 91.209 |  |  |  |  |  |  |
| Herr Frank virus 1 | 47.14 | 47.279 | 46.863 |  |  |  |  |  |
| Red-banded snake chuvirus | 45.921 | 45.967 | 45.738 | 46.688 |  |  |  |  |
| Wenling-fish chuvirus | 42.166 | 42.127 | 41.935 | 43.299 | 42.824 |  |  |  |
| Hardyhead chuvirus | 41.98 | 42.117 | 41.705 | 40.552 | 42.088 | 38.302 |  |  |
| Sanxia atyid shrimp virus 4 | 37.816 | 37.861 | 37.173 | 37.42 | 36.476 | 33.731 | 33.805 |  |

The amino acid pairwise identities of the G and N proteins between two marine chelonian chuviruses and the freshwater chelonian chuvirus was 85.9 and 83.4%, respectively; whereas the pairwise identity between marine chelonian chuviruses was 99.6 and 100%, respectively. The L and G proteins are the same length between freshwater chelonian chuvirus (STCV-1) and marine chelonian chuviruses (KTCV-1 and LTCV-1), whereas the SCTV-1 N protein is 2 amino acids smaller than the marine chelonian chuviruses. The highest percent amino acid identity for the G protein of chelonian chuviruses to other piscichuviruses was 35.7–36.7% to Guangdong red-banded snake chuvirus-like virus (GenBank accession numbers: MG600009) (Table 1 and supplemental data). The highest percent amino acid identity for the chelonian chuvirus N protein to other piscichuviruses was 37.7–38.4% to Guangdong red-banded snake chuvirus-like virus (GenBank accession numbers: MG600009) (Table 1 and supplemental data).

#### Phylogenetic analysis of *Chuviridae*

To understand the relationship of these chelonian chuviruses, putatively classified as Piscichuvirus testudinae, to other chuviruses; phylogenetic analyses using the amino acid sequences of each predicted protein were performed. Phylogenetic analysis of the predicted L protein amino acid sequences from 58 chuviruses using Maximum-Likelihood analysis demonstrated that STCV-1 KTCV-1, and LTCV-1, clustered with other piscichuviruses, and this genus had a branch length of 0.7037 from other chuviruses. All three turtle chuviruses clustered together in the putative Piscichuvirus testudinae species, with a branch length of 0.6286 and bootstrap value of 100%. The P. testudinae species shared a most recent common ancestor with *Piscichuvirus franki* (branch length of 0.7283) with a 24% bootstrap value. (Fig. 4).

**Figure 4.**
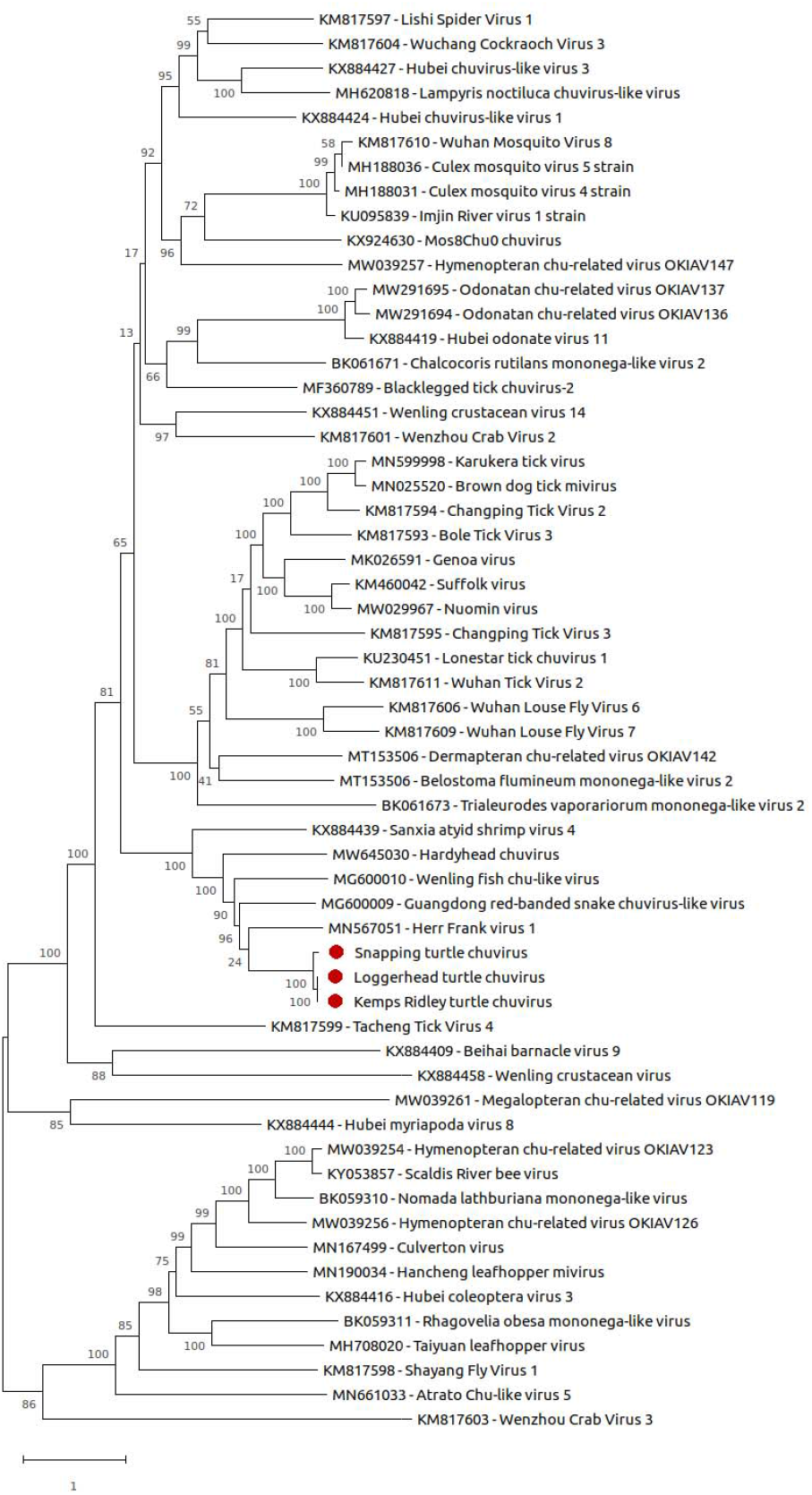

Of note, when running multiple sequence alignments for the phylogenetic analysis, it was noted that after the N coding region there is a fourth possible open reading frame (ORF) in piscichuviruses via NCBI ORF finder. This fourth ORF is consistent with what is annotated as VP4 or hypothetical protein in other chuviruses that have circular genomes, such as Wuhan tick virus 2 (GenBank accession No. MZ965027), Suffolk virus (GenBank accession No. NC028403), and lonestar tick chuvirus (GenBank accession No. NC030204). Within the genus *Piscichuvirus* that was previously deposited on GenBank, this coding region is 225–276 bases (74–91 amino acids) long.

#### *In situ* identification of a novel chuvirus using RNAscope™assay

To determine if the turtle chuviruses colocalized within the inflamed central nervous system (CNS) tissue, RNAscope™ *in situ* hybridization (ISH) was performed. In the alligator snapping turtle, the ISH demonstrated disseminated, strong, punctate reactivity for STCV-1 RNA in areas of inflammation throughout various areas of the central nervous system, predominantly the grey matter (Fig. 1). In the cerebellum, small neurons of the internal granular layer had strong intracytoplasmic reactivity (Fig. 1b). In the brainstem, there was strong to moderate intracytoplasmic reactivity in neurons with central chromatolysis. Strong reactivity was disseminated throughout all layers of the olfactory bulb, predominantly in the granular cell layer (data not shown). The cerebrum had strong, but less distributed probe signal, as compared to other brain sections. The optic tectum had disseminated, strong, intracytoplasmic ISH reactivity throughout the grey matter (Fig. 1d). The spinal cord had multifocal, strong reactivity in neuronal cytoplasm and around nuclei of glial cells. Other tissues were also tested for STCV-1 nucleic acid; however, non-CNS tissues from this alligator snapping turtle, and the brain tissue from an alligator snapping turtle without encephalitis lacked STCV-1 nucleic acid signal.

For KTCV-1, the viral RNA specific probe showed disseminated, strong reactivity in areas of inflammation and degeneration throughout various areas of the Kemp’s ridley turtle central nervous system. Similar to the STCV ISH, signal predominantly located within the grey matter (Fig 2). In the brainstem, there was strong to moderate intracytoplasmic reactivity in large and small neurons, and ependymal cells (Fig 2b). In the midbrain, strong reactivity was within the neuronal cytoplasm and around nuclei of glial cells (data not shown). The cerebrum had strong reactivity that was disseminated throughout the tissue section, predominantly within the gray matter and ependyma (data not shown). There was no ISH reactivity against viral RNA within the olfactory bulb (data not shown). However, there was mild ISH reactivity throughout the leptomeninx. The spinal cord had a few multifocal, strong ISH-positive puncta in the cytoplasm of neurons (Fig. 2d). Similar to STCV-1, other tissues were tested for KTCV-1 nucleic acid, including non-CNS tissues from this Kemp’s ridley turtle, brain tissue from a non-encephalitic Kemp’s ridley turtle, and brain tissue from a Kemp’s ridley turtle with bacterial meningitis. All of these tissues lacked reactivity for KTCV-1 nucleic acid.

Additionally, the STCV-1 probe and the KTCV-1 probe were also tested for cross-reactivity (e.g., STCV-1 tested on KTCV-1 infected Kemps ridley turtle). Both probes failed to detect the other virus (e.g., STCV-1 probe failed to detect KTCV-1).

Due to the high percent nucleotide (99%) identity between KTCV-1 and LTCV-1 probe regions, it was predicted that the KTCV-1-based probe would detect LTCV-1 RNA within the tissues of the loggerhead turtle. The KTCV-1 probe showed multifocal ISH reactivity in various areas of inflammation and vacuolation of central nervous system, e.g., optic tectum, cranial nerve, cerebrum, and cerebellum (Fig. 3). In the optic tectum, there was strong, disseminated intracytoplasmic reactivity within neurons, small neurons, and glial cells in all tissue layers (data not shown). The cranial nerve had fine puncta of the ISH signal widely disseminated throughout tissue section with rare aggregation (data not shown). In the cerebellum, there was disseminated, strong, intracytoplasmic ISH signal in small neurons and glial cells throughout the grey matter (Fig. 3b). The cerebrum had multifocal, strong ISH reactivity in the cytoplasm of ependymal cells, neurons, and glial cells disseminated throughout white, grey matter, and adjacent leptomeninx; where there were associated lymphocytic infiltrates (Fig. 3d). No ISH reactivity was observed in the olfactory bulbs.

#### Viral isolation

Samples from the alligator snapping turtle and Kemp’s ridley turtle were inoculated onto chelonian cell lines and were tested for replication of STCV-1 and KTCV-1 via reverse transcription PCR (RT-PCR). Nucleic acid of STCV-1 was detected in P1 inocula of YBSLHt and SSTLu. Lysate from the spinal cord and CSF of the Kemp’s ridley turtle were both tested via RT-PCR. Only P1 inocula from the spinal cord origin of DBTLu, GTSp, and SSTLu were RT-PCR positive.

## Discussion

*Chuviridae* is the only family of single-stranded RNA viruses in which the genomic structures can be linear or circular, and can be non-segmented or segmented within the same family. The unique genomic feature of this viral family is believed to be transitional evidence of viral evolution between monopartite (*Mononegavirales*) and polypartite (*Bunyavirales and Orthomyxoviridae*) viral families (21). Currently, there are 14 genera of chuviruses such as *Mivirus* spp. (most common hosts: chelicerate), *Chuvivirus* spp. (hosts: crustacean), and *Piscichuvirus* spp. (most common hosts: vertebrate). Similar to other piscichuviruses, these turtle piscichuviruses were single-stranded RNA with linear genomic structure.

Based on the recent classification criteria of order *Jingchuvirales* (21), percent pairwise amino acid identity and phylogenetic tree of L amino acid sequences indicated that STCV-1, KTCV-1, and LTCV-1 all belong to a novel species and share a common ancestor with other piscichuviruses. However, as with most viral families, and especially for a newly discovered family such as *Chuviridae*, such classification criteria are likely subject to change. For example, the viruses were detected from animals with shared ecosystems (marine KTCV-1 and LTCV-1) have a relatively high polymerase percent identity (99.4%) as compared to alignments between viruses from non-overlapping ecosystems (e.g., KTCV-1 vs. STCV-1 = 91.6% and LTCV-1 vs STCV-1 = 91.2%). While sea turtles and alligator snapping turtles are found in different ecosystems, there is connectivity between their habitats through river systems. Sea turtles are known to forage within tidal areas of rivers (32) and sea turtle stranding networks occasionally document carcasses of freshwater chelonians, including alligator snapping turtles, in estuarine and marine areas that presumably originated from river outflows (Stacy, pers observation). In addition, all host species examined in this study share the most recent common ancestor and are classified under *Americhelydia* clade, which is comprised of chelydroids (snapping turtles) and chelonioids (sea turtles, mud turtles, and hickatee) (33). As more information is gained about the diversity of chuviruses and their host restrictions through future studies, it is foreseeable that the currently proposed Piscichuvirus testudinae ultimately may be divided into two or more species (e.g., freshwater [chelydoid] and marine [chelonioid]).

Furthermore, the phylogenetic analysis included previously deposited sequences and it was noted that some previously characterized viruses have percent amino acid identities that are only slightly above the genera cutoff, e.g., Lishi spider virus 1 (38.5% amino acid identity to Wuchang cockroach virus 3), Sanxia atyid shrimp virus 4 (37.3% amino acid identity to Herr Frank virus-1), Wēnlǐng crustacean virus 14 (33.3% amino acid identity to Wenzhou crab virus 2) (see Table 1 and supplemental data). Notably, Sanxia Atyid shrimp virus 4, is the only piscivirus whose host is not a vertebrate. Given the host difference and how close it is to the numerical percent identity cutoff (31%) (21), other criteria, such as genetic distances and biological properties (e.g., vertebrate vs. invertebrate host), might be considered as additional criteria for future re-classification of the order *Jingchuvirales* after more of these viruses have been characterized.

For the genetic comparison of all chelonian chuviruses, the open reading frames of the predicted *L* and *G* genes of turtle chuviruses were slightly larger than other vertebrate chuviruses. However, the open reading frame of *N* gene was smaller than other vertebrate chuviruses, except Wēnlǐng fish chu-like virus. The open reading frames of predicted viral protein 4 (VP4), that are not commonly annotated in other chuviruses, were annotated in all chelonian chuviruses. The size of these VP4 regions were 318 bases long in all chelonian chuviruses with 100% amino acid pairwise identity between marine turtle chuviruses and 77.4% between freshwater and marine turtle chuviruses. Interestingly, both termini of all chelonian chuviral genomes were complimentary (with 2–3 mismatches). Additionally, intergenic regions were TA-rich (64.7%). Both of these features can lead to the formation of the inverted repeat and secondary structure, i.e., hairpin loop and Internal Ribosome Entry Sites (IRES)(34). The formation of these secondary structures benefits of host-virus interaction, i.e., viral RNA synthesis, splicing (34, 35), and host genome integration (36).

Even though the *Chuviridae* family was recently discovered through metagenomic sequencing in 2015, the timescale phylogenetics and EVEs suggest that chuviruses have co-evolved with a broad host species and have circulated in mosquitoes for approximately 190 million years (20, 25, 29). The number of known chuviruses has gradually increased within the past 5 years, reflecting the abundance of undiscovered chuviruses in various animals, but primarily within arthropods. ICTV, therefore, formed a chuvirus taxonomic working group for the future expansion of this family (21). However, the in-depth evolution and clade classification of chuviruses have not been well established due to the limited availability of variable genomic regions. Similar to other viruses, species identification of chuviruses mostly relies on the large gene, which encodes for the highly conserved viral polymerase (21). Finer classification of viruses, i.e., genotyping and evaluating temporospatial relationships, utilizes more variable genomic regions, such as the hemagglutinin for morbillivirus and influenza virus genotyping (37, 38). Currently, 144 sequences of chuviral G gene are available on GenBank (assessed on August 3, 2022). However, only 50 chuviral G gene sequences are from different ecological events. Therefore, the G gene or complete genome of chuviruses are needed to inform evolution timescale and the nucleic acid/amino acid substitution rate in future studies.

To date, chuviruses have not been definitively associated with disease in animals. The first potential association was the identification of Herr Frank virus 1 in snakes that were individually diagnosed with ulcerative stomatitis, osteomyelitis, or cloacitis (22). However, these snakes were also co-infected with reptarenavirus and hartmanivirus and no tissue localization studies were performed, so it is not possible to attribute disease to the chuvirus infection. Subsequently Nuomin virus was isolated from the serum of febrile human patients (30), but again, *in situ* studies were lacking and a clinically silent infection could not be ruled out. This current study identified the first *in situ* evidence of chuviral disease, in three different chelonians from two different ecosystems. In this study, all cases had severe predominantly lymphocytic meningoencephalomyelitis manifesting as severe clinical deficits that precipitated death or humane euthanasia.

The colocalization of RNAscope^™^ signal chuviral RNA and areas of inflammation and necrosis support the pathogenicity of these novel chuviruses. Thus, for the first time, the pathogenic impact of chuviruses has been demonstrated in vertebrates. Chuviruses should be considered in cases of idiopathic encephalitis, especially in chelonids. The identification of closely related chuviruses in other reptiles and fishes suggests that chuviruses should be considered in those species as well. Further surveillance is required to better determine the impact of chuviruses on these, and other animals. Because of the poor sequence similarity between known vertebrate chuviruses, random sequencing will likely be required in any vertebrate host other than those from which chuviruses have already been detected.

All of the chelonian host species in which chuvirus was found are considered imperiled as reflected in their classification as threatened or critically endangered by the International Union for Conservation of Nature and current (sea turtles) or proposed (alligator snapping turtle) listing under the US Endangered Species Act of 1973. Affected turtles included mature adults, which are especially vital to chelonian population stability and recovery (39). Notably, both the alligator snapping turtle and the loggerhead turtle were in relatively good nutritional condition at the time of death and did not have any apparent underlying condition suspected to have predisposed to viral infection. The potential to infect and cause disease in relatively healthy individuals represents a significant wildlife health concern. In addition, the alligator snapping turtle was believed to have been transported and released outside of its range, which raises the possibility of human-mediated pathogen pollution. Future studies are needed to understand the diversity and prevalence of chuvirus among chelonians, pathogenesis of infections, transmission pathway(s), and host-virus interaction.

## Material and methods

### Gross necropsy and sample collection

All animals died spontaneously or ultimately were euthanized (using pentobarbital solutions) due to advanced morbidity or persistent neurological abnormalities that were unresponsive to treatment. Gross necropsy included systematic evaluation of all organ systems. Sexual maturity was determined by evaluation of the reproductive system. Fresh tissue samples, including the brain, spinal cord, and CSF (Kemp’s ridley only) were aseptically collected and stored at −80°C until thawed for nucleic acid extraction. Fixed samples were preserved in neutral-buffered 10% formalin for histopathology.

### Histopathology

After 24 to 48 hours of formalin fixation, formalin-fixed tissues were serially dehydrated and embedded in paraffin. Formalin-fixed paraffin embedded (FFPE) tissue blocks of samples were sectioned at 5 μm onto glass slides and stained with hematoxylin & eosin (H&E). Selected tissues also were stained using the Ziehl-Neelsen acid fast method.

### Alligator snapping turtle and loggerhead turtle: RNA extraction and viral enrichment

Preserved cerebrum of an alligator snapping turtle was removed from RNAlater™(Thermo Fisher Scientific) and tapped on Kimwipes™(Kimberly-Clark) to remove excessive RNAlater™solution before proceeding with RNA extraction. Cerebrum from the alligator snapping turtle and brainstem from the loggerhead turtle were homogenized in 450 μL 1× phosphate-buffered saline (PBS) by using a TissueLyser LT (Qiagen) at 35 Hz for 2 minutes with a sterile stainless steel 0.5 mm metal bead (Qiagen). Homogenized samples underwent depletion of host and bacterial ribosomal RNA using a modified previously published protocol (40). Briefly, homogenates were centrifuged at 17,000 × *g*, at room temperature for 3 minutes to remove large cellular debris and bacteria. Supernatants were collected and passed through an 0.8 μm PES filter (Sartorius) for the removal of smaller debris and bacteria. Filtrates were treated with a cocktail of 2.0 μL of micrococcal nuclease and 1.0 μL benzonase (2,000,000 gel units/mL Micrococcal nuclease, NEB Inc, and >250 units/μL Benzonase™nuclease, Millipore Sigma) in 7.0 μL of resolving enzyme buffer to remove any free nucleic acids before performing the nucleic acid extraction (41). Trizol™LS Reagent (Thermo Fisher Scientific) was used for RNA extraction following the manufacturer’s protocol.

### Alligator snapping turtle and loggerhead turtle: Random MinION sequencing

The MinION sequencing library was prepared using a random strand-switching protocol as previously published (42). In short, the purified RNA concentration of the brain and spinal cord of the alligator snapping turtle and loggerhead turtle was quantified by using Qubit RNA HS assay kit (Qubit 3.0 fluorometer; Thermo Fisher Scientific) and reverse transcribed by using the SuperScript IV reverse transcriptase kit (Thermo Fisher Scientific) with 1 μM PCR-RH-RT primer (5’ -ACTTGCCTGTCGCTCTATCTTC<u>NNNNNN</u>-3’) as a reverse primer and 10 μM strand-switching oligo (5’ -TTTCTGTTGGTGCTGATATTGCTGCCATTACGGCCmGmGmG-3’; both synthesized by Integrated DNA Technologies [IDT]) for a strand switching reaction. The reaction mixture was incubated at 50°C for 30 minutes, 42°C for 10 minutes, and 80°C for 10 minutes before being bead purified at a 0.7× beads:solution ratio (KAPA Biosystem). cDNA then was used as a template for barcoding PCR following ONT’s protocol (SQK-LSK110 with EXP-PBC096) and LongAmp *Taq* 2× Master Mix (NEB, Ipswich, MA). The barcoded amplicons were bead purified at a 0.8× beads:solution ratio before being pooled by equal volume with libraries from unrelated samples and a library generated from HeLa RNA (ThermoFisher)

(negative control library). The library pool was used for end repair and ligation of the sequencing adapter using suggested kits for ligation sequencing amplicons (SQK-LSK110) library preparation. Bead purification was done after end repair and adapter ligation at 1.0× and 0.4× beads:solution ratios, respectively. The library was washed with long fragment buffer (LFB), and eluted in elution buffer (EB) before mixing with sequencing buffer (SBII) and loading beads II (LBII), following ONT’s protocol. The final library was then loaded onto a FLO-MIN106 R9.4.1 flow cell in a MinION Mk1B sequencing device (ONT).

Sequencing was controlled using MinKNOW v.21.06.0 (ONT) for 29 hours with default settings and hourly MUX scans. Post-sequencing analysis workflow followed the randomly primed, MinION-based sequencing as previously described (42, 43). In brief, raw reads were basecalled, with the GPU version of Guppy v.6.1.3 (ONT) using the following command: guppybasecaller -i <input path> -s <save path> --flowcell <FLO-MIN106> --kit <SQK-LSK110> --device auto --calib_detect. Porechop (http://githup.com/rrwick/Porechop) was used for trimming and demultiplexing using the following command: porechop -i <input reads> -b <output directory> --require_two_barcodes --adapter_threshold 99 --extra_end_trim 0 --check_reads 1000000 > output_file.txt. Potential viral reads were screened using all reads by pairwise aligning to non-redundant Basic Local Alignment Search Tool (BLAST+) using BLASTn database (updated on June 04, 2022) through the Georgia Advanced Computing Research Center (GACRC) using default settings.

Reference-based mapping was also used to identify chuviral reads within the sample. Hence, reads were first filtered by alignment using a custom Centrifuge index. Centrifuge v.1.0.4. was used following the previously described protocol using a viral database (42, 44), with the addition of chuviral sequences and the green sea turtle genome (GCF_015237465.1_rCheMyd.pri.v2, accessed in July 2021; closest, most complete genome to alligator snapping turtle). Reads that failed to align within this Centrifuge analysis were then filtered for bacterial reads using a publicly available Centrifuge index for bacteria and archaea (p_compressed_2018_4_15.tar.gz) (https://ccb.jhu.edu/software/centrifuge/manual.shtml) (44). Reads that were greater than 50 nt and aligned to chuvirus in the first Centrifuge analysis or reads that failed to align to the host, other viruses, or bacteria were mapped to Wēnlǐng fish chuvirus-like virus (GenBank accession No. MG600011) in Geneious Prime 2019.1.3 (https://www.geneious.com) (map to reference setting with 15 iterations and medium sensitivity). The coding regions of both STCV-1 and LTCV-1 were annotated by using the NCBI ORF finder (https://www.ncbi.nlm.nih.gov/orffinder/) and by comparing to the annotations of other piscichuviruses on Geneious Prime 2019.1.3. Gaps of consensus sequence were filled using targeted MinION sequencing

### Kemp’s ridley turtle: RNA extraction and random Illumina sequencing

RNA was extracted from the brainstem tissue sample using an RNeasy Mini Kit (Qiagen) according to the manufacturer’s instructions. cDNA library was generated using a NEBNext Ultra RNA Library Prep Kit (Illumina) and sequenced on the iSeq 100 Sequencing System. Raw data (10,782,756 paired-end reads) were processed to remove host reads by first running Kraken v2 (45) (https://ccb.jhu.edu/software/kraken2/) against a custom database created using the green sea turtle genome (GenBank assembly accession: GCA_000344595, accessed in March 2020). *De novo* assembly of the remaining paired-end reads (742,979) was performed using

SPAdes v3.15.3 with default parameters. The assembled contigs were then subjected to BLASTX searches in OmicsBox v2.0 (BioBam Bioinformatics) against the National Center for Biotechnology Information (NCBI) nonredundant protein database. The coding regions of both KTCV-1 were annotated by using the NCBI ORF finder and by comparing to the annotations of other piscichuviruses. Gaps of consensus sequence were filled using additional PCR reactions for Sanger sequencing.

### Probe design and RNAscope™ *in situ* hybridization (ISH) assay

FFPE blocks from the encephalitic alligator snapping turtle, encephalitic Kemp’s ridley turtle, and encephalitic loggerhead turtle were sectioned at 5 μm on ColorFrost Plus™glass slides (Erie Scientific) using new microtome blades between samples to avoid cross contamination. FFPE blocks of cerebrum and cerebellum from an unrelated alligator snapping turtle that lacked encephalitis, unrelated Kemp’s ridley turtles (e.g., a Kemp’s ridley turtle without encephalitis and a Kemp’s ridley turtle with bacterial meningitis), and unrelated loggerhead turtles (a loggerhead turtle without encephalitis and loggerhead turtles with lymphohistiocytic meningoencephalitis) were included in this study as a negative tissue control. RNAscope ™2.5 HD double z-probes (Catalog No. 1138851-C1, Advanced Cell Diagnostics, Inc.) were designed using the L gene of the STCV-1 genomic consensus sequence from random, deep MinION sequencing at bases 1375-2368, and the L gene of KTCV-1 genomic consensus sequence from random, deep Illumina sequencing at bases 1535-2585. Due to the limited availability of reference genes in alligator snapping turtles and Kemp’s ridley turtle, the genome of the common snapping turtle (*Chelydra serpentina*) (GenBank accession No. ML689093) was used in the probe design to minimize non-specific probe binding to host tissues during hybridization. A probe targeting *Bacillus subtilis* dihydrodipicolinate reductase (DapB) gene was used as a negative probe control. A uniquely conserved area of ribosomal RNA of chelonian species (green turtle, loggerhead turtle, and common snapping turtle) was selected to design the probe and was used as a positive control to validate the RNA integrity of the samples (Catalog No. 1231491-C1, Advanced Cell Diagnostics, Inc). RNAscope™ISH was performed by following manufacturer’s protocol for RNAscope™2.5 HD detection reagent – RED (document number 322360-USM and 322452-USM). Briefly, all tissue sections were incubated at 60°C and deparaffinized using freshly prepared xylene and ethanol. Antigen retrieval was performed with hydrogen peroxide (10 minutes at room temperature), heat (15 minutes at 100°C), and protease enzyme (30 minutes at 40°C) using the provided kit reagents. Subsequently, probes were hybridized under following condition: probe hybridization at 40°C for 2 hours, AMP1 hybridization at 40°C for 30 minutes, AMP2 hybridization at 40°C for 15 minutes, AMP3 hybridization at 40°C for 30 minutes, AMP4 hybridization at 40°C for 15 minutes, AMP5 hybridization at room temperature for 30 minutes, and AMP6 hybridization at room temperature for 15 minutes. Slides were washed with 1× wash buffer (ACD Bio) at room temperature for 2 minutes after each hybridization. Ultimately, slides were counterstained with hematoxylin, cover slipped, and dried overnight prior to evaluation with light microscopy.

### Phylogenetic Analysis

The *L*, *G*, and *N* open reading frames from the alligator snapper turtle and Kemp’s ridley chuviruses, with 55 other complete viral nucleotide sequences under order *Jingchuvirales* from GenBank (Fig. 4), were translated using Geneious. All alignments and phylogenetic analyses were conducted in MEGA X. Multiple sequence alignments of each coding region were done separately with ClustalW and MUSCLE’s default setting. Best substitution models of aligned amino acid sequences for *L, G* and *N* were selected based on the lowest Bayesian information criterion (BIC) and Akaike scores using the best substitution model analysis for maximum-likelihood analysis in MEGA X. Based on the best substitution model analysis, phylogenetic analysis for *L* amino acid coding sequences was constructed using the Maximum-Likelihood (ML) method and Le Gascuel matrix (LG) + observed amino acid frequencies (F) + 5 discrete gamma categories distribution (G) + invariant sites (I) substitution model with 500 bootstrap replicates. Subtree-Pruning-Regrafting level 3 was used for ML tree inference. Phylogenetic analysis for *G* amino acid coding sequence was constructed using Maximum-Likelihood method and Whelan and Goldman (WAG) + amino acid frequencies model (F) model + 5 discrete gamma categories distribution (G) with 500 bootstrap replicates. Subtree-Pruning-Regrafting level 3 was used for ML tree inference. Phylogenetic tree for *N* amino acid coding sequences was constructed using Maximum-Likelihood method and General Reverse Transcriptase (rtREV) + amino acid frequencies model (F) + 5 discrete gamma categories (G) substitution model at 500 bootstrap replicates. Subtree-Pruning-Regrafting level 3 was used for ML tree inference. All gaps and missing data were used to construct all phylogenetic trees in this study.

### Viral isolation

To isolate novel chuviruses, alligator snapping turtle brain, and Kemp’s ridley turtle spinal cord and CSF were inoculated onto 4 chelonian cell lines. These cell lines included gopher tortoise spleen (GTSp), yellow belly slider heart (YBSHt), diamondback terrapin lung (DBTLu), and softshell turtle lung (SSTLu) established from *Gopherus polyphemus, Trachemys scripta, Malaclemys terrapin*, and *Apalone ferox*, respectively. All cell lines were maintained in 32° C incubators in a humidified, 5% CO_2_ atmosphere. Cells were grown in T25 flasks using Minimum Essential Medium with Earle’s Balanced Salts, L-Glutamine (MEM/EBSS; GenClone), 10% heat inactivated fetal bovine serum (FBS; GenClone), nonessential amino acids (Caisson), penicillin-streptomycin solution (GenClone), amphotericin B (HyClone), and gentamicin (GenClone). Cell monolayers of GTSp, YBSHt, and DBTLu were grown until a confluency of 70-95% was reached. Cells of SSTLu did not reach full confluency, and were considered ready when clusters of cells covered approximately 10% of the flask. For all cell lines immediately prior to inoculation, media was removed and the flask was washed twice using sterile phosphate buffered saline (PBS).

To prepare tissues for P0 inoculation, brain and spinal cord were washed with PBS and finely minced using sterile scalpel blades. The minced tissue was mixed with 200 μl of completed medium, and further ground using a Pellet Pestles™(Fisher) and a microcentrifuge tube. Finally, ground tissue was mixed with 1800 μl of completed medium. For inoculations of spinal fluid, 500 μl of spinal fluid was mixed with 1500 μl of completed medium and mixed via pipet 5 times. A flask of each cell line was inoculated with 500 μl of each inoculum and incubated for 60 minutes at room temperature with gentle rocking every 10 minutes. For mock inoculations, flasks were given 500 μl untreated completed media. Complete culture medium (4 mL) was added to each flask after the initial adsorption period, and returned to a 32° C, humidified 5% CO2 atmosphere.

To inoculate P1 flasks, 500 μl of P0 cell lysate was inoculated onto flasks of the same cell line using the protocol above. If P0 flasks showed cloudy media indicative of bacterial growth, prior to inoculation the P0 lysate was first centrifuged at 800 ×*g* for 30 seconds and the resulting supernatant was passed through a 0.22 μm inline syringe filter prior to inoculation.

Confirmation of viral growth in P1 lysate was performed by using a reverse transcription PCR (RT-PCR). Viral primers were designed to STCV-1 and KTCV-1 using Geneious. Mitochondrial cytochrome oxidase subunit 1 long was used as a reference gene control (46). All passages of CPE positive and negative lysates, and mock-infected lysates were tested for viral propagation. TRIzol™ LS was used to extract the RNA following the manufacturer’s suggested protocol. Reverse transcription was performed on extracted RNA, along with no-template controls and no-enzyme controls, using SuperScript™ IV first-strand synthesis system for RT-PCR (Invitrogen) with random hexamers (50 ng/μL). Subsequently, cDNA was amplified for STCV-1 (DreamTaq Green PCR master mix, 2X; Thermo Fisher) with the following thermocycyling conditions: 10μM of each primer set (STCV-1 FWD: AATTCAGGGTGGTGCAGGAG-3, and SCTV-1 REV: 5’ GCCACTCTCCTCGTTTCACA-3), 95°C for 1 min; 35 cycles of 95°C for 30 s, 53°C for 30 s, 72°C for 1min; 72°C for 5 min. For KTCV-1, similar thermocycling conditions were used KTCV-1 FWD: 5’GTTCTCAGACCGCGTTACCA-3 and KTCV-1 REV: 5’ CTCATTGGCGCATCAAGTCG-3) with 54°C annealing temperature. Bands were visualized with electrophoresis in a 1.5% agarose gel. Amplicons were purified (QIAquick^®^ PCR purification kit; Qiagen) following the manufacturer’s protocol and eluted in 30 μL of nuclease-free water (Qiagen). Final concentration and purity were measured (Qubit dsDNA HS assay kit, Qubit 3.0 fluorometer; Thermo Fisher Scientific). The purified PCR products and 5 μM of each primer were submitted to Eurofins Genomics (Eurofin Genomics LLC, KY) for bidirectional Sanger sequencing.

## Acknowledgement

We thank Dr. Paula Baker and staff and volunteers of the ARK and the UF-CVM Zoological Medicine service for their contributions to veterinary care and husbandry. Funding for cell line generation was provided by Morris Animal Foundation grant D18ZO-319. We are grateful to the participants in the Sea Turtle Stranding and Salvage Network (STSSN) for response to and documentation of stranded sea turtles. We also thank Nicole Stacy at the University of Florida for the cytological analysis and interpretation of cerebrospinal fluid. We declare no conflicts of interest with respect to the research, authorship, and/or publication of this article.

## Indices

Please see the attached files

