## Supplementary material for "Piscichuviral encephalitis in marine and freshwater chelonians: first evidence of jingchuviral disease": Caption

**Captions**

**Figure 1: Meningoencephalomyelitis, Alligator snapping turtle (*Macrochelys* sp.)**. (1a) Cerebellum: Lymphoplasmacytic perivascular cuffs (solid arrow) and infiltrates are widely disseminated in the grey matter and the adjacent leptomeninges (HE). (1b) A replicate section of the same tissue of Fig. 1a. There is strong ISH signal against snapping turtle chuvirus (STCV-1) in the cytoplasm of small neurons and glial cells throughout the grey matter and associated with the lymphoplasmacytic infiltrates. (1c) Optic tectum: Several neurons have central chromatolysis. Hematoxylin and eosin (HE). (1d) A replicate section of the same tissue of Fig.1c. There is intense ISH signal against STCV-1 within the neuronal and glial cytoplasm. The higher magnification in the bottom right corner shows the ISH signal within the neuronal cytoplasm.

**Figure 2**: **Meningoencephalomyelitis, Kemp's ridley turtle (*Lepidochelys kempii*).** (2a) Brainstem: Lymphocytic perivascular cuffs (solid arrows) and infiltrates are multifocally disseminated in the grey matter (HE). (2b) A replicate section of the same tissue of Fig. 2a. There is strong ISH signal against kemp’s ridley turtle chuvirus (KTCV-1) throughout the ependymal layer and within the cytoplasm of glial cells, small neurons, and large neurons of the grey matter adjacent to perivascular cuffs. (2c) Spinal cord: There are multifocal lymphocytic perivascular cuffs and vacuolation within the spinal cord parenchyma. Eosinophilic inclusion-like material is within the neuronal nuclei (solid arrows) (HE). (2d) A replicate section of the same tissue of Fig. 2c. Rare, scattered cells have strong, punctate, intracytoplasmic ISH signal.

**Figure 3: Meningoencephalomyelitis, Loggerhead turtle (*Caretta caretta*)**. (3a) Midbrain: Lymphocytic perivascular cuffs (solid arrows) and infiltrates are widely disseminated in the grey matter (HE). (3b) A replicate section of the same tissue of Fig.3a. There is abundant, intense ISH signal using the KTCV-1 probe within the neuronal and glial cytoplasm adjacent to the areas of lymphocytic perivascular cuffs. (3c) Cerebrum: Predominantly lymphocytic perivascular cuffs and infiltrates are multifocally disseminated in the grey matter and adjacent leptomeninx (HE). (3d) A replicate section of the same tissue of Fig.3a. There are numerous puncta of ISH signal using the KTCV-1 probe in glial cells associated with the areas of inflammation in the grey matter and leptomeninx.

**Table 1: Percent amino acid identities of predicted L amino acid sequences of viruses in *Piscichuvirus* genus.**

Table excerpts percent amino acid identities from the supplemental data.

*GenBank Accession number of piscichuviruses; LTCV-1: TBD, KTCV-1: TBD, STCV-1: TBD, Herr Frank virus 1: MN567051, Red-banded snake chuvirus: MG600009, Wenling-fish chuvirus: MG600010, Hardyhead chuvirus: MW645030, and Sanxia atyid shrimp virus 4: KX884439.

**Figure 4**: **Phylogenetic analysis of *Chuviridae***. **L amino acid sequences.** Phylogenetic analysis of complete L protein amino acid sequences of viruses representing each species in *Jingchuvirales* order. Complete L protein amino acid sequences were aligned by ClustalW and refined by MUSCLE with default settings. The phylogenetic analysis was performed on MEGA X using the Maximum-likelihood method and Le Gascuel matrix (LG) + observed amino acid frequencies (F) + 5 discrete gamma categories distribution (G) with parameter of 1.0728 + invariant sites (I) with 0.65% sites. substitution model with 500 bootstrap replicates. The tree with the highest log likelihood (-242567.81) is shown. The tree is drawn to scale, with bootstrap values and branch lengths measured in the number of substitutions per site. This analysis included 58 amino acid sequences. The chelonian chuvirus sequences of this study are denoted with red circles.
